# Interpretable attention model in transcription factor binding site prediction with deep neural networks

**DOI:** 10.1101/648691

**Authors:** Chen Chen, Jie Hou, Xiaowen Shi, Hua Yang, James A. Birchler, Jianlin Cheng

## Abstract

Due to the complexity of the biological factors that may influence the binding of transcription factors to DNA sequences, prediction of the potential binding sites remains a difficult task in computational biology. The attention mechanism in deep learning has shown its capability to learn from input features with long-range dependencies. Until now, no study has applied this mechanism in deep neural network models with input data from massively parallel sequencing. In this study, we aim to build a model for binding site prediction with the combination of attention mechanism and traditional deep learning techniques, including convolutional neural networks and recurrent neural networks. The performance of our methods is evaluated on the ENCODE-DREAM in vivo Transcription Factor Binding Site Prediction Challenge datasets.

The benchmark shows that our implementation with attention mechanism (called DeepGRN) improves the performance of the deep learning models. Our model achieves better performance in at least 9 of 13 targets than any of the methods participated in the DREAM challenge. Visualization of the attention weights extracted from the trained models reveals how those weights shift when binding signal peaks move along the genomic sequence, which can interpret how the predictions are made. Case studies show that the attention mechanism helps to extract useful features by focusing on regions that are critical to successful prediction while ignoring the irrelevant signals from the input.

## 1 Introduction

Transcription factors are proteins that bind to specific genomic sequences and affects numerous activities such as division, growth, development, and death. They regulate the rates of transcriptional activities of downstream genes through such binding events, thus acting as activators or repressors in the gene regulatory networks by controlling the expression level and the protein abundance of their targeted genes (Hobert, 2008). Chromatin immunoprecipitation-sequencing (ChIP-Seq) has been widely used to determine the interactions of a transcription factor and all its potential binding regions on genomic sequences. However, ChIP-Seq experiments sometimes require experimental materials that are infeasible to acquire, such as antibodies targeting specific transcription factors of interest. Thus, predictions of potential binding sites through computational methods are considered as possible alternative solutions. In addition, prediction of binding sites of transcription factors would facilitate many biological studies by providing resources as reference for experimental validation.

Many algorithms have been developed to solve this problem, including supervised learning and unsupervised learning methods. Unsupervised methods such as hidden Markov models (Mathelier and Wasserman, 2013; Mehta, et al., 2011) and hierarchical mixture models (Pique-Regi, et al., 2011) usually classify input genomic regions into different clusters by their affinity to certain transcription factors. Supervised learning approaches require ground truth during model training and aim to precisely predict the bound/unbound status of transcription factors in specific regions, including support vector machines (Djordjevic, et al., 2003; Zhou, et al., 2015), discriminative maximum conditional likelihood (Keilwagen, et al., 2019) and random forest (Hooghe, et al., 2012; Xiao and Segal, 2009). Both types of methods usually rely on prior knowledge about sequence preference data, such as position weight matrix (Sherwood, et al., 2014), which need additional steps to generate the required features. In addition, these features may be less reliable when no prior knowledge exists and inference based methods (such as de-novo motif discovery) are used to generate the input features (Keilwagen, et al., 2019).

More recently, methods based on deep neural networks (DNNs), such as DeepBind, TFImpute, and DeepSEA, have shown performances superior to traditional models (Alipanahi, et al., 2015; Zeng, et al., 2016; Zhou and Troyanskaya, 2015). Compared with the traditional methods, deep learning models have their advantages at learning high-level features from data with extremely large size. This property makes them ideal for the binding site prediction task since a huge amount of training data can be generated from the ChIP-Seq experiment data. Unlike many existing models that rely heavily on the quality of the input data and labor-intensive feature engineering, deep learning requires less domain knowledge or data pre-processing and is more powerful when there is little or no prior knowledge of potential binding regions.

Current studies in the protein binding site prediction tasks usually involve the combination of two deep learning architectures: convolutional neural networks (CNN) and recurrent neural networks (RNN). The CNN has the potential to extract both local features such as different genomic signals and regions (Kalkatawi, et al., 2019), while the RNN is better at learning useful information with long-term dependencies. Several popular methods for binding prediction, such as DanQ (Quang and Xie, 2016), DeeperBind (Hassanzadeh and Wang, 2016) and FactorNet (Quang and Xie, 2019), are built on such model architecture.

Recently, the concept of attention mechanism has achieved great success in neural machine translation (Luong, et al., 2015) and sentiment analysis (Wang, et al., 2016). It enhances the ability of DNNs by focusing on the information that is highly valuable to successful prediction within the input. Combining with RNNs, it allows models to learn the high-level representations of input sequences with long-range dependencies. In addition, it makes such models more interpretable by assigning attention weights to different positions of the input regarding their importance. Different visualization methods have been developed to explore the relationship between the input and output sequences using the attention mechanism, including the alignment view (Alkhouli and Ney, 2017) and the extra layer of interpretability (Xu, et al., 2015). With these techniques, we expect that introducing the attention mechanism to binding site prediction would increase prediction accuracy as well as the level of interpretability for existing CNN-RNN architecture models.

In this study, we propose a transcription factor binding prediction tool called DeepGRN, which is a combination of attention mechanism and DNNs. We show that this combination improves the performance of the existing CNN-RNN architectures. Also, we explored the attention weights extracted from the trained networks to explain how the input data are utilized through the attention mechanism.

## 2 Methods

### 2.1 Datasets from ENCODE-DREAM Challenge

The datasets used for model training and benchmarking are from the 2016 ENCODE-DREAM *in vivo* Transcription Factor Binding Site Prediction Challenge. The detailed description of the pre-processing of the data can be found at https://www.synapse.org/#!Synapse:syn6131484/.

Information about the binding status of the transcription factors is generated from ChIP-Seq experiments and used as ground truth. Chromatin accessibility information (DNase-Seq data), and RNA-Seq data are provided as input features for model training. We use the similar organization of input features introduced by FactorNet (Quang and Xie, 2019): DNA Primary sequence, Chromatin accessibility information (DNase-Seq data) are transformed into sequential features, while gene expression and annotations are transformed into non-sequential features.

#### 2.1.1 Transcription factor binding data

Transcription factor binding data from ChIP-Seq experiments is the target for our prediction. The whole genome is divided into bins of 200bp with a sliding step size of 50bp (i.e., 250-450bp, 300-500bp). Each bin falls into one of the three types: bound, unbound, and ambiguous. Bins overlapping with peaks and passing the Irreproducible Discovery Rate (IDR) check with a threshold of 5% (Li, et al., 2011) are labeled as bound. Bins that overlap with peaks but fail to pass the reproducibility threshold are labeled as ambiguous. All other bins are labeled as unbound. We do not use any ambiguous bins during training or validation process according to the common practice. Therefore, each bin in the genomic sequence will either be a positive site (bounded) or a negative site (unbounded) in our labels.

#### 2.1.2 DNA primary sequence

Human genome release hg19/GRCh37 is used as the reference genome. We expand each bin by 400bp in both its upstream and downstream, resulting in a 1000bp region around it. The sequence of this region is represented by a 1000×4 bit matrix by 1-hot encoding with each row represented a nucleotide. In addition, sequence uniqueness (also known as “mappability”) plays an important part in the quality for Next-Generation Sequencing (NGS) since low mappability sequences may introduce a bias in sequencing or its analysis (Sholtis and Noonan, 2010), Duke 35bp uniqueness score is included as an extra input feature. Scores ranging from 0 to 1 are assigned to each position as the inverse of occurrences of a sequence with the exceptions that the scores of unique sequences are 1 and scores of sequences occurring more than four times are 0 (Derrien, et al., 2012). As a result, the sequence uniqueness is represented by a 1000×1 vector for each input bin. The ENCODE Project Consortium has provided a blacklist of genomic regions that produce artifact signals in NGS experiments (ENCODE Project Consortium, 2012). We exclude input bins overlapping with these regions from training data and set their prediction scores to 0 automatically if they are in target regions of prediction.

#### 2.1.3 DNase-Seq data

Chromatin accessibility reflects the accessibility of regions on a chromosome and is highly correlated with transcription factor binding events (Pique-Regi, et al., 2011). DNase-Seq can be used to obtain genome-wide maps of chromatin accessibility information as chromatin accessible regions are usually more sensitive to the endonuclease DNase-I than non-accessible regions (Madrigal and Krajewski, 2012). DNase-Seq results for all cell types are provided in the Challenge datasets in the BigWig format. Normalized 1x coverage score is generated from the BAM files using deepTools (Ramirez, et al., 2014) with bin size = 1 and is represented by a 1000×1 vector for each input bin.

#### 2.1.4 Gene expression and annotation

The annotation feature for each bin is encoded as a binary vector of length 6, with each value represent if there is an overlap between the input bin and each of the six genomic features (coding regions, intron, promoter, 5’/3’-UTR, and CpG island). The annotations file for these features is provided by the FactorNet Repository (https://github.com/uci-cbcl/FactorNet/tree/master/resources). Also, RNA-Seq data can be used to represent the differences in gene expression levels among different cell types. The principal component analysis is performed on the Transcripts per Million (TPM) normalized counts from RNA-Seq data of all cell types provided by the Challenge. The first eight principal components of a cell type are used as expression scores for all bins from that cell type, generating a vector of length 8. Both of the non-sequential features are fused into the first dense layer in the deep learning model.

### 2.2 Training, validation and evaluation

In this study, training, validation, and testing of our models are performed on the 12 different transcription factors officially used by the DREAM Challenge. The scope of the Challenge is restricted to chromosomes 1-22 and X. All these chromosomes except for the held-out chromosomes (chromosome 1, 8 and 21) of the cell types for training (Table S1) are used to train the models. The labels on chromosome 11 are excluded during the training phase and used as validation data for hyperparameter tuning.

Among all transcription factors, eight of them have one cell type (called “leaderboard cell type”) for additional model tunning. Labels of the leaderboard cell types were not released during the challenge. Instead, participants can submit predictions and query their performances on Chromosomes 1 and 21 of the leaderboard cell types for at most ten times in order to improve their models. Under this rule, for transcription factors with a leaderboard cell type, we use the top 10 models with the highest auPRC from validation for a re-validation on Chromosomes 1 and 21 of the leaderboard cell types. The models with the best performance in this procedure are picked for the final benchmark. For transcription factors without any leaderboard cell type, the models with the highest auPRC on chromosome 11 are picked for the final benchmark (Table S1, Figure S1).

Due to the sparsity of the binding events on the whole genome, the positive labels are generally much less than the negative labels (i.e., transcription factor CTCF has less than 0.35% of positive labels in training data), resulting in an extremely unbalanced classification problem. To effectively evaluate the performance of different models, auROC (area under the curve of recall versus the false positive rate) and auPRC (area under the curve of precision versus the recall) are used as scoring metrics. The auPRC metric generally performs better than auROC and accuracy score when negative labels are predominant, as it does not consider the number of true negatives in evaluation (Saito and Rehmsmeier, 2015). For this reason, hyperparameters are tuned based on the auPRC. We also include the recalls at different FDR levels as an additional metric. Recalls at fixed FDR levels measure the ability of the classifier to identify true positives when too many false positives are not allowed.

### 2.3 Deep neural network models with attention mechanism

The shape of each sequential input is L×(4+1+1) for each region with length L after combining all sequential features (DNA sequence, sequence uniqueness, and Chromatin accessibility in Section 2.1). Sequential inputs are generated for both the forward strand and the reverse complement strand. The weights in all layers of the model are shared between both inputs to form a “Siamese” architecture (Mueller and Thyagarajan, 2016; Qin and Feng, 2017; Quang and Xie, 2019). For each transcription factor, the two output values from the forward and reverse inputs are merged at the end of the model by taking the maximum or mean of those values depending on the auPRC from validation. Vectors of non-sequential features from gene expression data and genomic annotations are fused into the model at the first dense layer. The overall architecture of our model is shown in Figure 1a.

**Figure 1.**
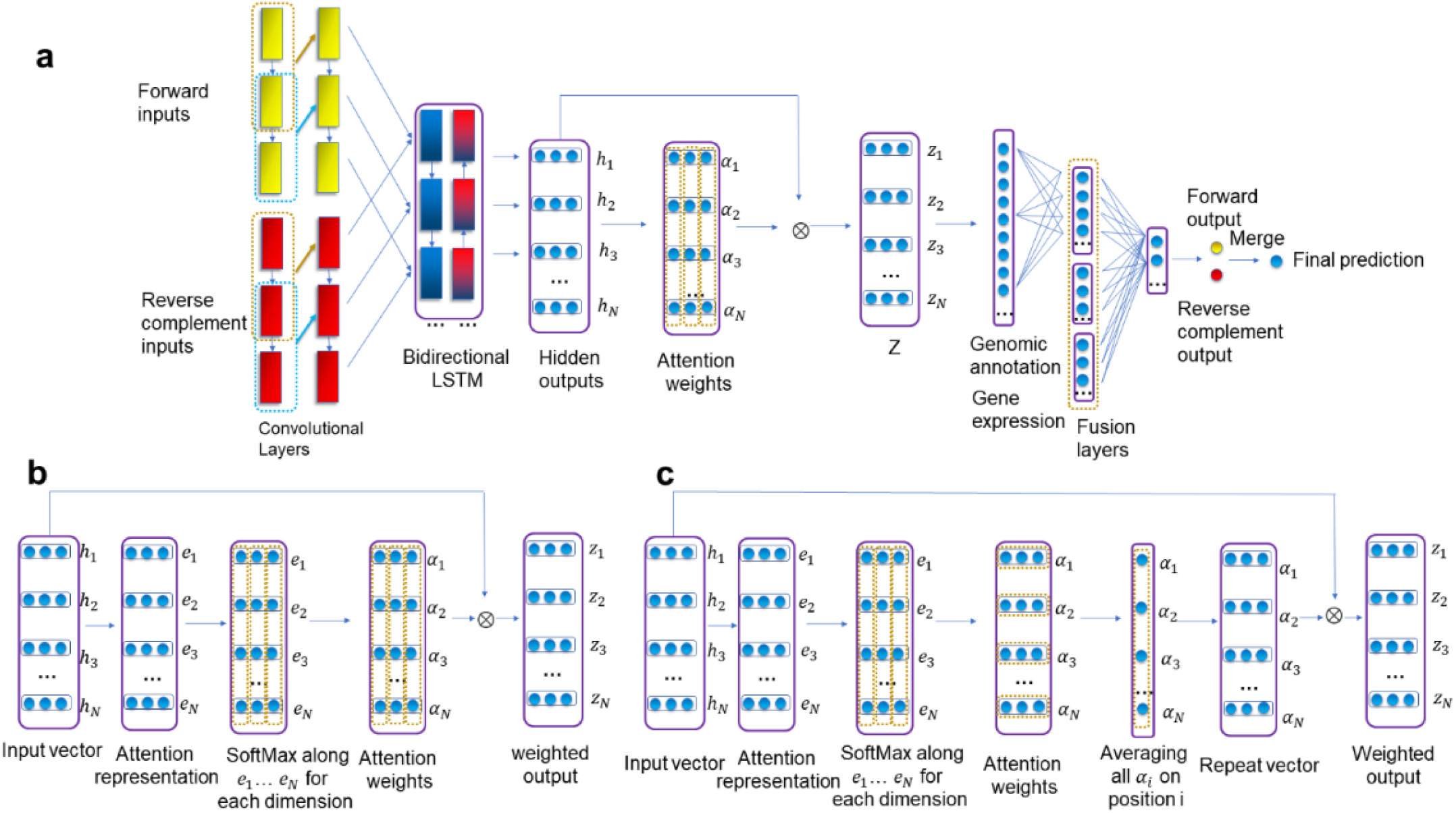
The diagram of the deep neural network architecture. **(a)** Convolutional and bidirectional LSTM (BiLSTM) layers use both forward and reverse complement features as inputs. Attention weights are computed from hidden outputs of LSTM and then are used to compute the weighted representation Z through a Kronecker product. Z is flattened and fused with non-sequential features (genomic annotation and gene expression). The final score is computed through dense layers with sigmoid activation and merging of both forward and reverse complement inputs. **(b, c)** Details about the implementation of attention mechanism with or without merging the weights.

We use Long Short-term Memory (LSTM) nodes as recurrent units in our model. We regard all the hidden vectors H = [*h*_1_, *h*_2_,…, *h*_*N*_] from the LSTM layer as the representation of sequential features, where N is the length of the LSTM input. Each hidden vector *h*_*i*_ is an array with length R. The attention mechanism will compute attention weight vectors *α*_1_, *α*_2_,…, *α*_*N*_ and weighted output representation Z.

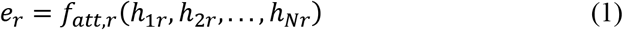

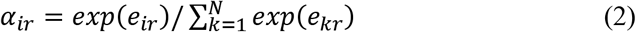

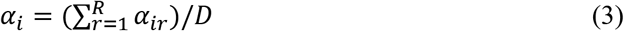

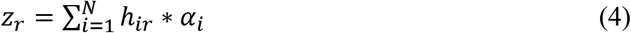

Here, *e*_*r*_ is the attention input representation for all LSTM output vectors at dimension *r*, which is learned from a multilayer perceptron *f*_*att, r*_. Attention weights *α*_*ir*_ are computed from the input representation at dimension *r* with Softmax normalization (Figure 1b). The dimension of the attention weights can be reduced from N × R to N×1 by averaging at each position (Figure 1c). Finally, the prediction scores are generated from dense layers with sigmoid activation function and merged from both forward and reverse complement inputs.

Typical hyperparameters for neural networks (learning rate, network depth, and dropout rates) as well as the hyperparameters specific to our model (the dimension of attention weights, merging function the two output scores) are tuned. The complete description of hyperparameters and their possible options are summarized in Table S3.

There are 51676736 bins in total on training chromosomes in the labels, resulting in 51676736×*n* potential training samples for each transcription factor, where *n* is the number of available cell types for training. Due to limited computing capacity, we use the iterative training process s with downsampling the negatives (Keilwagen, et al., 2019; Quang and Xie, 2019). In each epoch, we first sample *N*_*neg*_ negative bins from all negative labels. *N*_*neg*_ is proportional to the number of all positive bins, and the ratio of negative bins to positive ones tested are 1, 5, 10, and 15. These sampled negative bins are mixed with all positive bins for the training of the next epoch. To make the training process more effective, for transcription factors CTCF, FOXA1, HNF4A, MAX, REST and JUND, as they have a large number of positive labels, during each epoch we randomly sample one 200-bp region from each ChIP-Seq peak in the narrow peak data as positive instances for training. Adam (Kingma and Ba) is used as the optimizer, and binary cross-entropy is used as the loss function. The default number of epochs is set to 60, but the training will be early-stopped if there are no improvements in validation auPRC for five consecutive epochs.

The source code is available at https://github.com/jianlin-cheng/DeepGRN

## 3 Results

### 3.1 Overall benchmarking on evaluation data

We compared our results with four different algorithms from the top four teams in the final leaderboard of the ENCODE-DREAM Challenge (Table 1). The transcription factors, chromosomes, and cell types for evaluation are the same as those used for the final rankings. Since auPRC is better at capturing the difference of prediction performances with imbalanced labels than auROC and recalls at different FDR levels, we choose auPRC as our primary benchmarking metric. Based on auPRC, our attention model has better performance on 69.23% (9/13) of the prediction targets than Anchor (Li, et al., 2019), 69.23% (9/13) than FactorNet (Quang and Xie, 2019), 76.92% (10/13) than Catchitt (Keilwagen, et al., 2019), and 92.31% (12/13) than Cheburashka (Lando, et al., 2016). Among all the methods, our method achieved the highest auPRC on 6 targets: CTCF/induced pluripotent stem cell (iPSC), FOXA1/liver, FOXA2/liver, GABPA/liver, HNF4A/liver, and REST/liver.

**Table 1.**
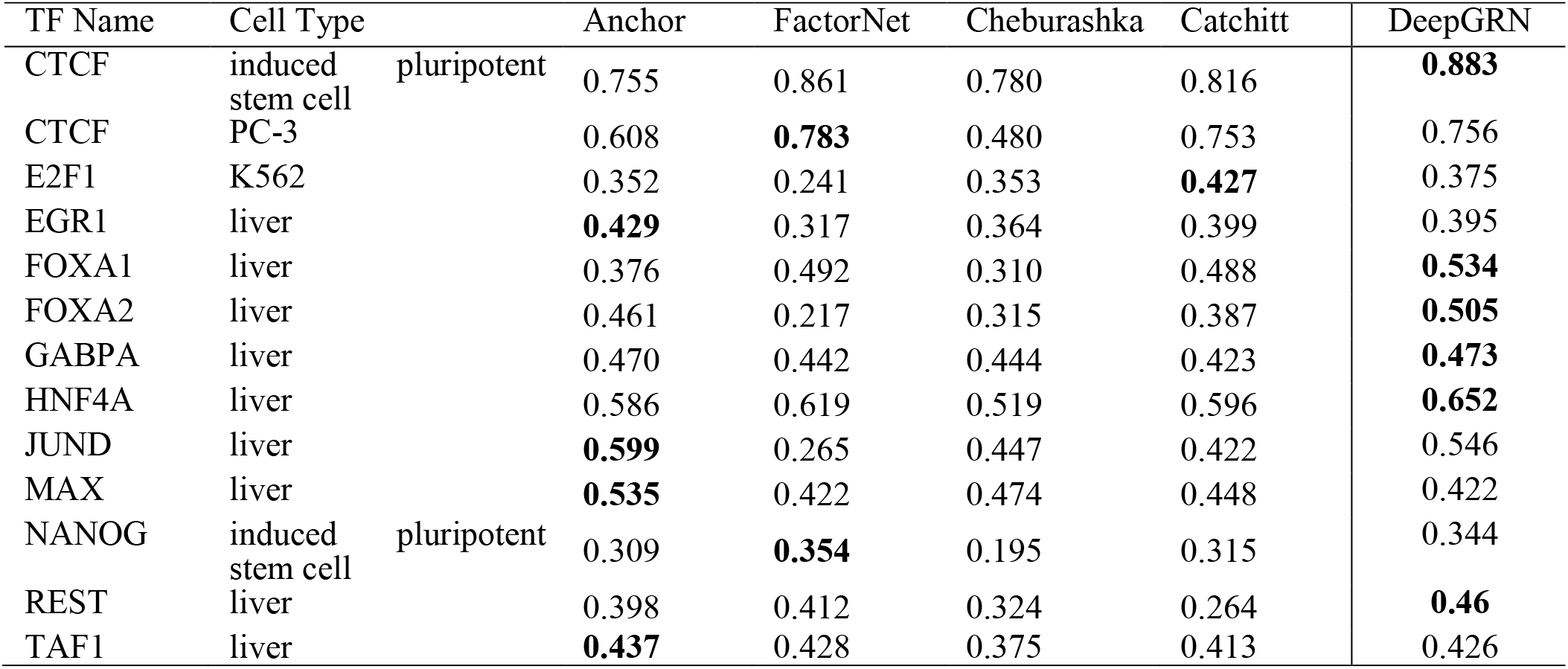
The results of our attention model and top four algorithms in the DREAM Challenge (Bold denotes the highest auPRC among all methods).

### 3.2 Performance improvements from attention mechanism

In addition to the comparisons with top 4 methods in the challenge, we also benchmarked the improvements from the attention mechanism in comparison with similar CNNs-RNNs model without attention on all test chromosomes (Table S2, S4). The model with attention achieved better or equal performance on 7 targets with all scoring metrics and 10 targets with 3 out of 4 metrics except for the recall at FDR level 0.1 (Figure 2a). The recall at 0.1 FDR is too stringent for the comparison, as its scores are less than 0.3 in 11 targets for both models. This makes it less reliable than other metrics. The percentage of improvements in average auROC across all targets is 0.66% (from 0.9812 to 0.9877). The improvement in average auPRC is 21.84% (from 0.4302 to 0.5208). The largest improvements for attention model in auPRC are achieved for JUND (0.262), FOXA2 (0.204), E2F1 (0.144), and EGR1 (0.118). For CTCF, the auPRC is slightly decreased by 0.027 in PC-3 cell but increased by 0.199 in iPSC.

**Figure 2.**
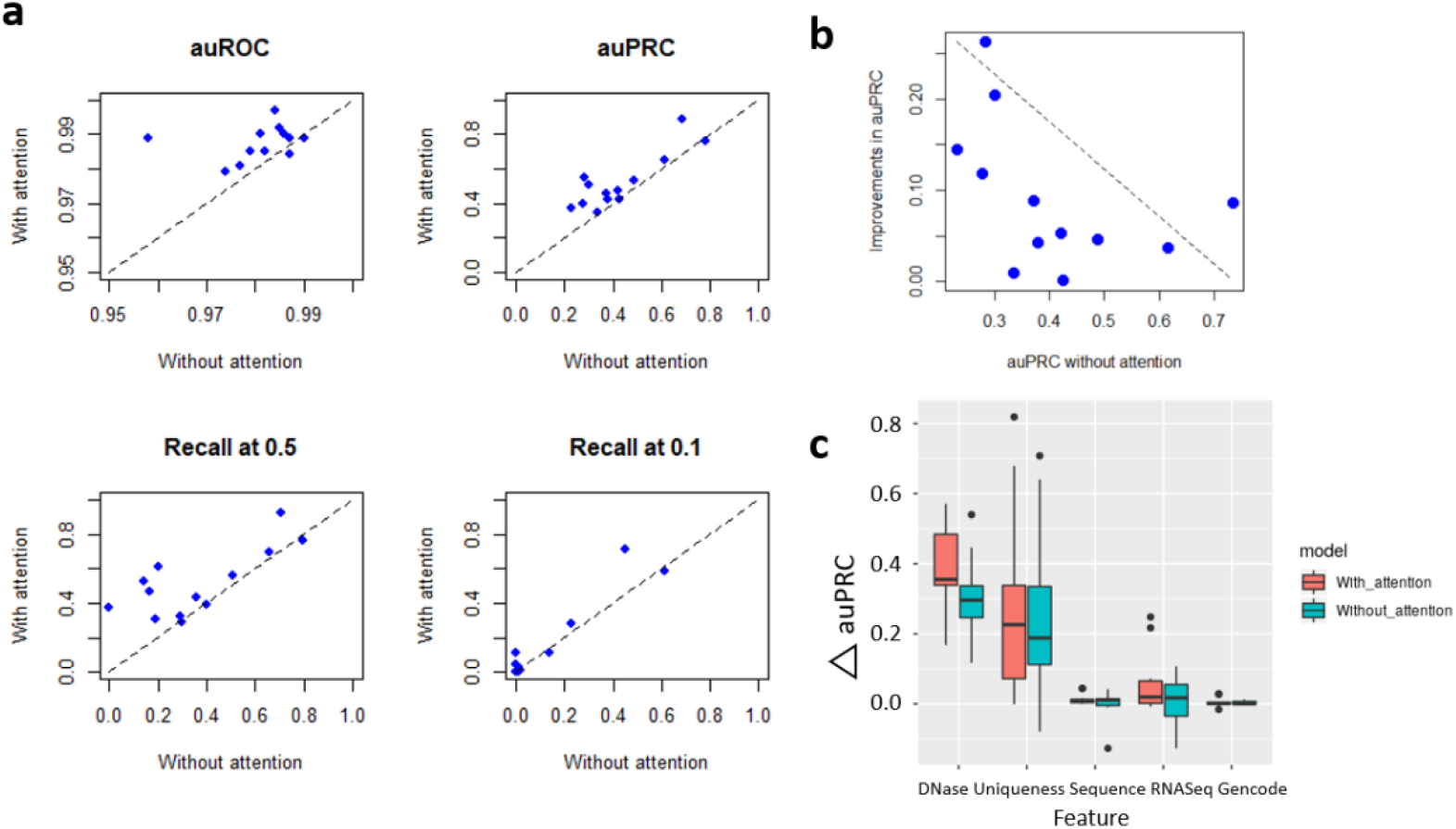
Performance improvements from attention mechanism. (**a**) Performance gain from attention mechanism for four metrics. (**b**) Relevance of the auPRC in model without attention and improvements in attention models. ρ: Pearson Correlation Coefficient, σ: Spearman Correlation Coefficient. (**c**) Change of auPRC after permutation of each feature in both two types of models.

We found that the transcription factors originally with low prediction accuracy have more improvements from the attention mechanism than those with relatively high prediction accuracy (Figure 2b). We also tested the difference between the two model types in terms of the importance of different features. To achieve this, we randomly permute each type of the sequential feature (DNase-Seq, sequence, or uniqueness) among bins, or randomly switch the order of the vector for a non-sequential feature (genomic elements or RNA-Seq) for each bin in the evaluation data and compute the auPRC from new predictions based on the permuted data (Figure 2c). The results show that models with attention mechanism are more susceptible to the permutation of DNase-Seq (*p* = 0.013), and the differences in for all other features are insignificant (*p* > 0.05) in Wilcoxon Signed-Rank Test, suggesting the main improvements of performance in attention models are from in a better match of the DNase-Seq peak signal.

### 3.3 Interpretation of attention mechanism

The visualization of attention weights is an important way to understand how the predictions are generated from the models. Since the input length of our model can be as long as 1000bp, the actual positions that lead to transcription factor binding events cannot be directly determined from the prediction result. To analyze the relationship between attention weights and the position of transcription factor binding events, we extract the attention weights from sequential input data on the forward strand and compare them with the ChIP-Seq fold change data. These fold change values are calculated as fold-enrichment of ChIP-Seq signal to the control at each nucleotide in the training chromosomes from the auxiliary data files of the ENCODE-DREAM Challenge in Bigwig format. These fold change signals at nucleotide level have the best representation of the location of transcription factor binding events. The coherence of these fold change values and the attention weights can be used to test if the attention mechanism has helped the models to focus on the important part of the sequence.

The trained model for transcription factor CTCF on four different cell types H1-hESC, HeLa-S3, HepG2, and K562 are used as examples (Figure 3). From the fold change results provided in the Challenge data, we located a genomic region from 276100 bp to 277550 bp on Chromosome 10 that has a clear ChIP-Seq peak signal. The attention weights of two input bins within this region are extracted from the model. For each cell type and genomic region, the plots in the first row are the fold change signals and the second row are the attention weights averaged on each position. To make the trend more obvious, we only choose top 10% dimensions with the highest variances from the weight vectors. Since the dimensions of attention weights are reduced by the convolution and pooling layers, their lengths are less than the fold change values. So, the plots are aligned on the X-axis to represent the relative position of fold change and averaged attention weights. We show that the averaged attention weights put more focus on the actual binding region for each cell type and the focusing point shifts along with the shift of transcription factor binding signals.

**Figure 3.**
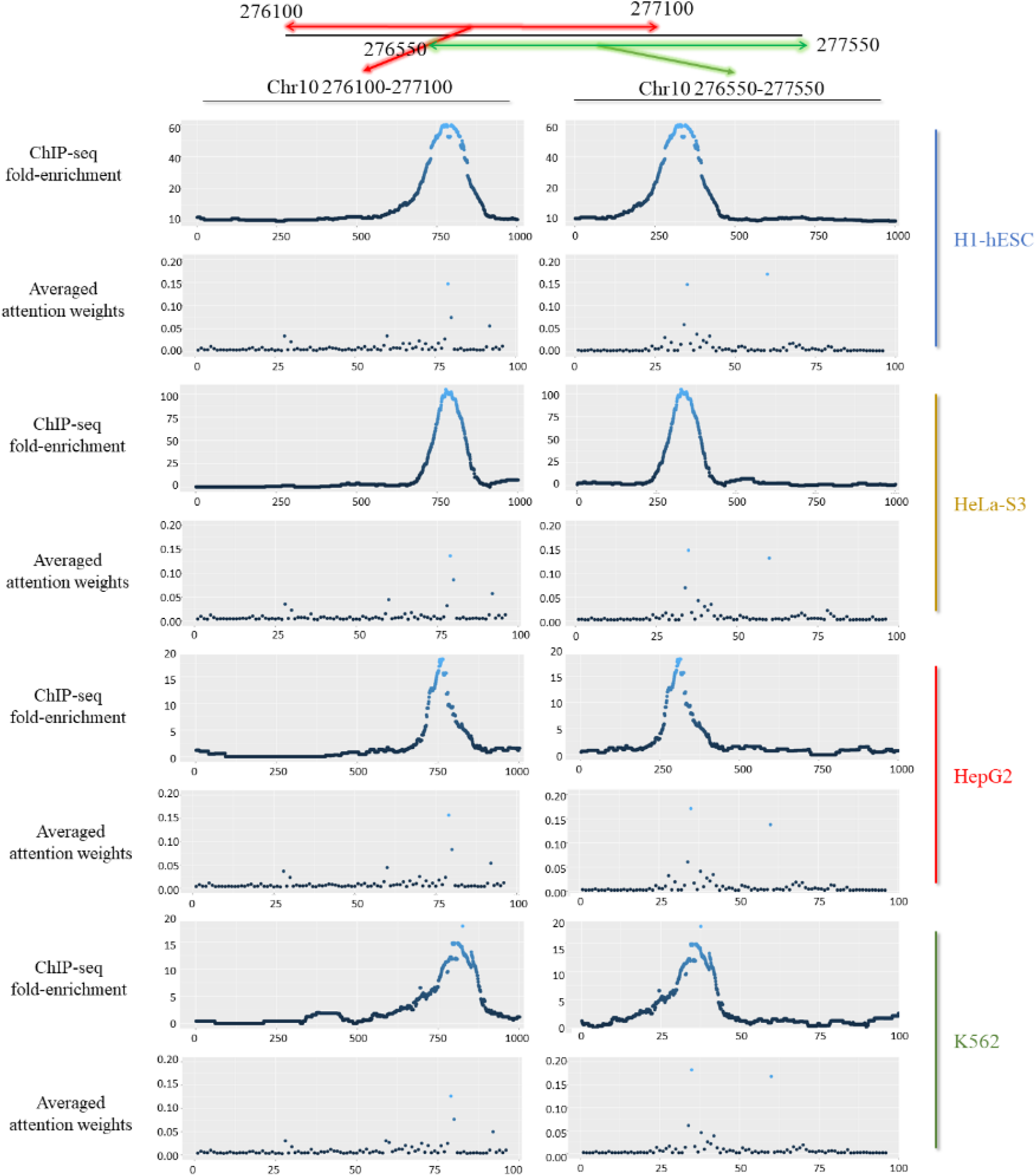
The relationship between ChIP-Seq peak and attention weights. Red and green lines denote two input genomic regions (Here we only choose from the positive strand). For each of four cell types (H1-bESC, HeLa-S3, HepG2, and K562), the plots in the first row represent the enrichment of fold changes in ChIP-Seq for the two regions and the plots in the second row represent the average of attention weights.

Also, to investigate how the attention mechanism focuses on local regions that play more important roles and discard the irrelevant signals that may cause interference, we choose transcription factor JUND in two cell types MCF-7 and HCT116 as examples. Both cell types have a positive ChIP-Seq peak within the genomic region from 179050 bp to 180050 bp on Chromosome 10. Since DNase-Seq data is the critical feature for prediction, we include the plots of DNase coverage score for the corresponding regions. Figure 4 shows the comparisons of the inputs in the same genomic region with the different distributions of DNase coverage scores. In either of the sample with one clear DNase peaks (MCF-7) or multiple ambiguous signals (HCT116), the learned attention weights put high scores on correct regions with true ChIP-Seq fold change peaks.

**Figure 4.**
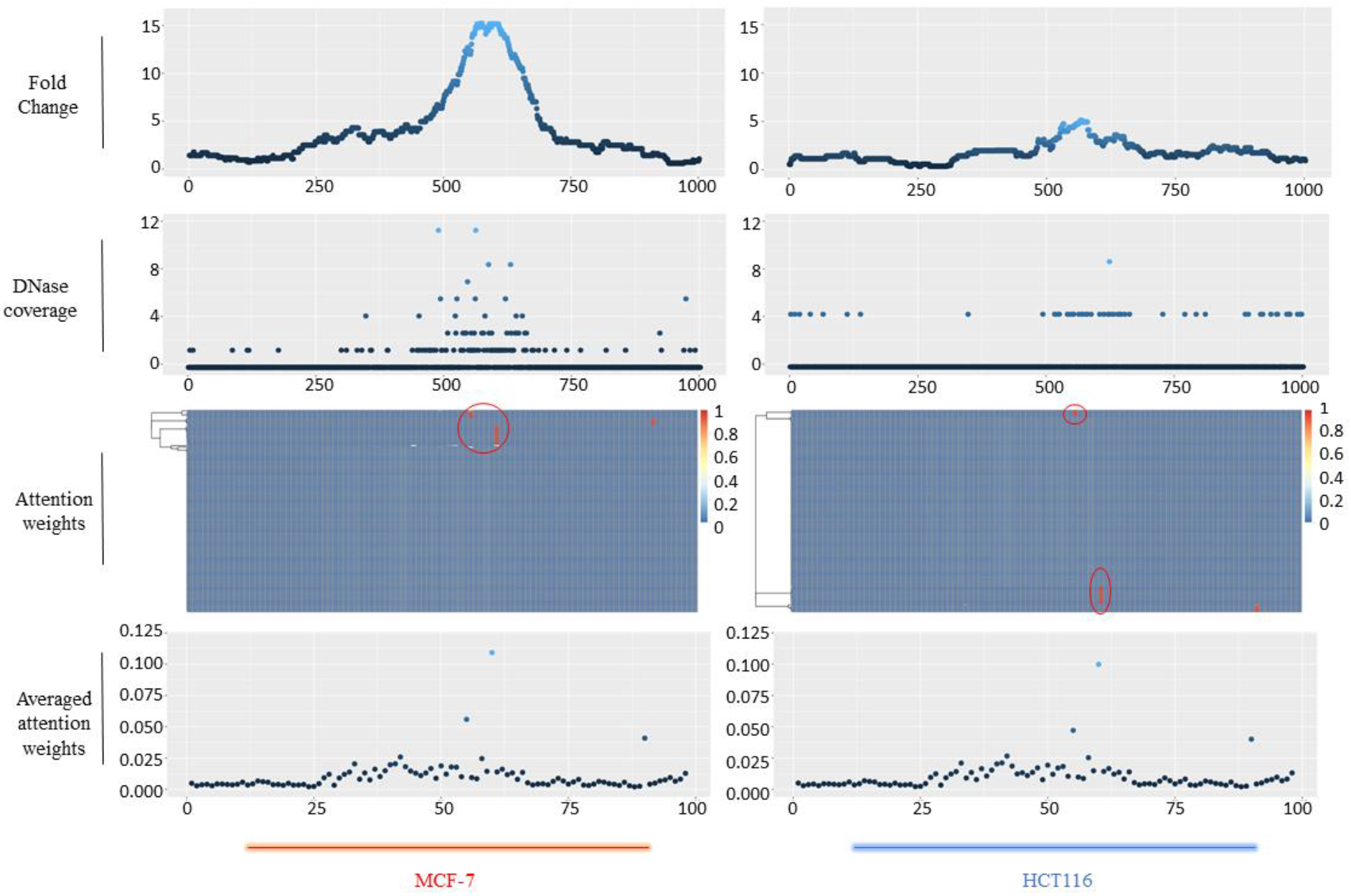
Visualization of attention weights and the ChIP-Seq/DNase peak on the corresponding region of two cell types (MCF-7 and HCT116) from 179050 bp to 180050 bp on Chromosome 10. From top to bottom are the trend of ChIP-Seq fold change signal, normalized 1x DNase coverage, heatmap representing the attention weights (*α* ∈ *R*^*d* × *l*^, where l is the length of the tensor in the attention layer and d is the dimension of the attention weights at each position), and the average attention weights at each position for the two cell types. The heatmap are clustered by rows for better visualization. Blocks highlighted by red circles are those weights with high scores in different dimensions and in concordance with the true ChIP-Seq fold change peaks.

To test the ability of attention weights to recognize motifs that contribute to binding events from the genomic sequences, we use an approach similar to DeepBind (Alipanahi, et al., 2015). We first acquired the coordinates on the relative positions of maximum attention weights from all positive bins in test datasets and extracted subsequences with a length of 20bp around these coordinates. Then we ran MEME (Bailey and Elkan, 1994) to detect *de novo* motifs from those subsequences and TOMTOM (Gupta, et al., 2007) to compare the discovered motifs with known motifs in the JASPAR database (Khan, et al., 2018). We show that these maximum attention weights do not come from the DNase-Seq peaks near the motif region by coincidence (Figure 5). The full list of all motifs identified from the subsequences by TOMTOM is provided in Table S5.

**Figure 5.**
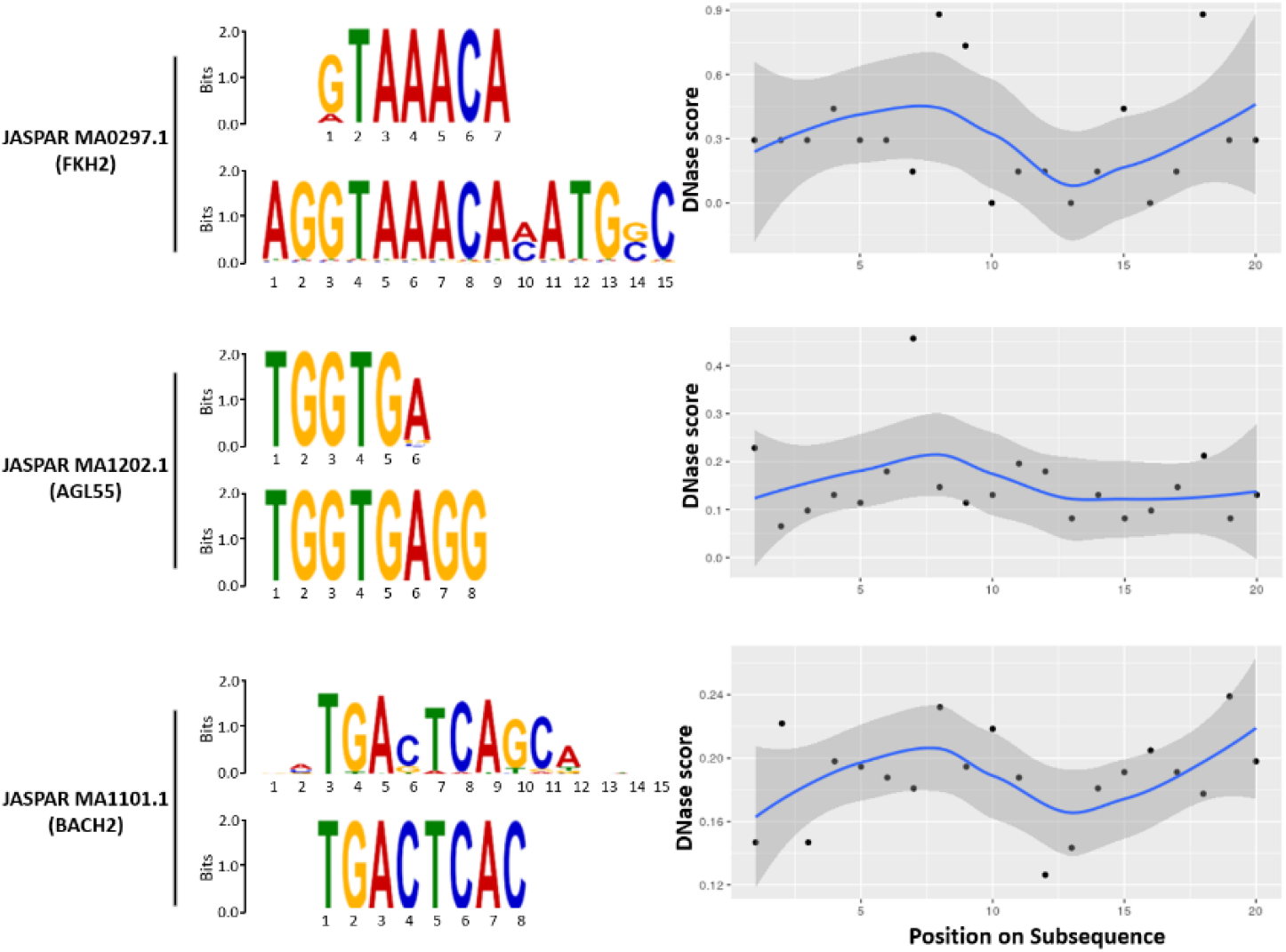
Comparisons of known motifs and matching motifs learned from attention mechanism. For each motif, the sequence logo in the top panel refers to a known motif in JASPAR database while the bottom panel denotes the matching motif detected by MEME from the subsequence near the position of the largest attention weight. The averaged DNase-Seq scores from all subsequences that match the corresponding motif are shown in the right.

## 4 Discussion

The attention mechanism is attractive in various machine learning studies and has achieved superior performance in image caption generation or natural language processing tasks (Vaswani, et al., 2017; Yang, et al., 2016). Recurrent neural network models with attention mechanism are particularly good at tasks with long-range dependency in input data. Inspired by these works, we introduce the attention mechanism to DNN models for transcription factor binding site prediction in order to improve their accuracy and interpret how the predictions are made.

We designed a new tool (DeepGRN) that incorporates the attention mechanism with the CNNs-RNNs based model by applying attention normalization before or after the LSTM layer. We tested our models on the ENCODE-DREAM Challenge datasets. The result shows that performances of our models are competitive with the top 4 methods in the Challenge leaderboard. Compared with similar deep learning models without attention mechanism, adding attention layers increases the prediction performance in almost all metrics adapted in the Challenge. It is also worth mentioning that the DNase-Seq scores are the most critical feature in the attention mechanism from our experiments. Many prediction tools for binding site prediction before the challenge, such as DeepBind or TFImpute, are not able to utilize the DNase-Seq data. Thus, they are not as suitable as the four methods that we used for benchmarking in this study.

Perhaps more importantly, the attention weights learned by the model provide a good way for interpretation, which has always been a challenging problem for deep learning applications in bioinformatics. Unlike many other machine learning methods such as regression or decision trees, the neural networks are often criticized for lack of interpretability. Nowadays, DNNs with moderate sizes can consist of millions of parameters, and it is extremely difficult to understand how the models utilize different parts of the input features. The attention weights provide a way to explore the dependencies between input and output. By comparing true ChIP-Seq fold change peaks with attention weights, we show how attention weights shift when the fold change peaks move along the DNA sequence. Our method also explains how the normalized scores are assigned to each part of the input feature according to their importance to successful prediction. Moreover, we demonstrate that the attention mechanism can precisely focus on the part of the input that is critical for the binding event while assigning lower scores to irrelevant regions with ambiguous DNase signals. This ability may explain the performance gained in our models.

Due to the rules of the DREAM Challenge, we only use very limited types of features in this work. However, if more types of data (such as sequence conservation or epigenetic modifications) are available, they can all be transformed into sequential formats and may further improve the prediction performance through the attention mechanism.

## Acknowledgements

We wish to thank the organizers of ENCODE-DREAM *in vivo* Transcription Factor Binding Site Prediction Challenge.

## Funding

This work has been supported by NSF grants (IOS1545780 and DBI1149224).

### Conflict of Interest

None declared.

