## Supplemental Table 3 and 4 for "Interpretable attention model in transcription factor binding site prediction with deep neural networks"

**Table S3** Hyperparameters selection for model training. Due to the limitation of computing capacity 50 sets of the combination of these hyperparameters are sampled without replacement for evaluating the configurations.

|  | Convolution layers | | | | LSTM layers | | | Attention layer | | | Dense Layers | | | |
| --- | --- | --- | --- | --- | --- | --- | --- | --- | --- | --- | --- | --- | --- | --- |
| Learning rate | Layers | Kernel Size | Dimension | Dropout rate | Layers | Dimension | Dropout rate | Position | *f_att_* | Dimension reduction | Layers | Dimension | Dropout rate | Merge |
| 0.01,  0.001,  0.0005,  0.0001 |  |  |  |  |  |  |  |  |  |  |  |  |  |  |
|  | 0,1,2 |  |  |  |  |  |  |  |  |  |  |  |  |  |
|  |  | 15,20,  30,34 |  |  |  |  |  |  |  |  |  |  |  |  |
|  |  |  | 64,128 |  |  |  |  |  |  |  |  |  |  |  |
|  |  |  |  | 0.1,0.5 |  |  |  |  |  |  |  |  |  |  |
|  |  |  |  |  | 1,2 |  |  |  |  |  |  |  |  |  |
|  |  |  |  |  |  | 32,64 |  |  |  |  |  |  |  |  |
|  |  |  |  |  |  |  | 0,0.1,0.5 |  |  |  |  |  |  |  |
|  |  |  |  |  |  |  |  | Before/  after LSTM |  |  |  |  |  |  |
|  |  |  |  |  |  |  |  |  | MLP, *F(x)=x* |  |  |  |  |  |
|  |  |  |  |  |  |  |  |  |  | Yes,No |  |  |  |  |
|  |  |  |  |  |  |  |  |  |  |  | 2,3,4 |  |  |  |
|  |  |  |  |  |  |  |  |  |  |  |  | 64,128 |  |  |
|  |  |  |  |  |  |  |  |  |  |  |  |  | 0.1,0.5 |  |
|  |  |  |  |  |  |  |  |  |  |  |  |  |  | Max,  Mean |

**Table S4** Types of trained models without attention mechanism. The models are from the trained best models of Factornet since we contrasted similar basic CNNs-RNNs architecture with it in this study.The models are downloaded form https://github.com/uci-cbcl/FactorNet/tree/master/models. For each transcription factor, we benchmarked all of the Factornet models in test datasets and pick the model of highest auPRC (model for transcription factor CTCF is chosen based on the average in two cell types) to represent the performance without attention mechanism.

| TF | cell | The model with highest auPRC |
| --- | --- | --- |
| CTCF | induced_pluripotent_stem_cell | meta_RNAseq_Unique35_DGF |
| CTCF | PC-3 | metaGENCODE_RNAseq_Unique35_DGF |
| E2F1 | K562 | GENCODE_Unique35_DGF |
| EGR1 | liver | onePeak_DGF |
| FOXA1 | liver | onePeak_DGF |
| FOXA2 | liver | onePeak_Unique35_DGF |
| GABPA | liver | meta_RNAseq_Unique35_DGF |
| HNF4A | liver | onePeak_Unique35_DGF |
| JUND | liver | meta_Unique35_DGF_2 |
| MAX | liver | onePeak_Unique35_DGF_2 |
| NANOG | induced_pluripotent_stem_cell | GENCODE_Unique35_DGF |
| REST | liver | GENCODE_Unique35_DGF |
| TAF1 | liver | GENCODE_Unique35_DGF |
