## Supplementary figures and images for "Interpretable attention model in transcription factor binding site prediction with deep neural networks"

### Supplemental Figure 1

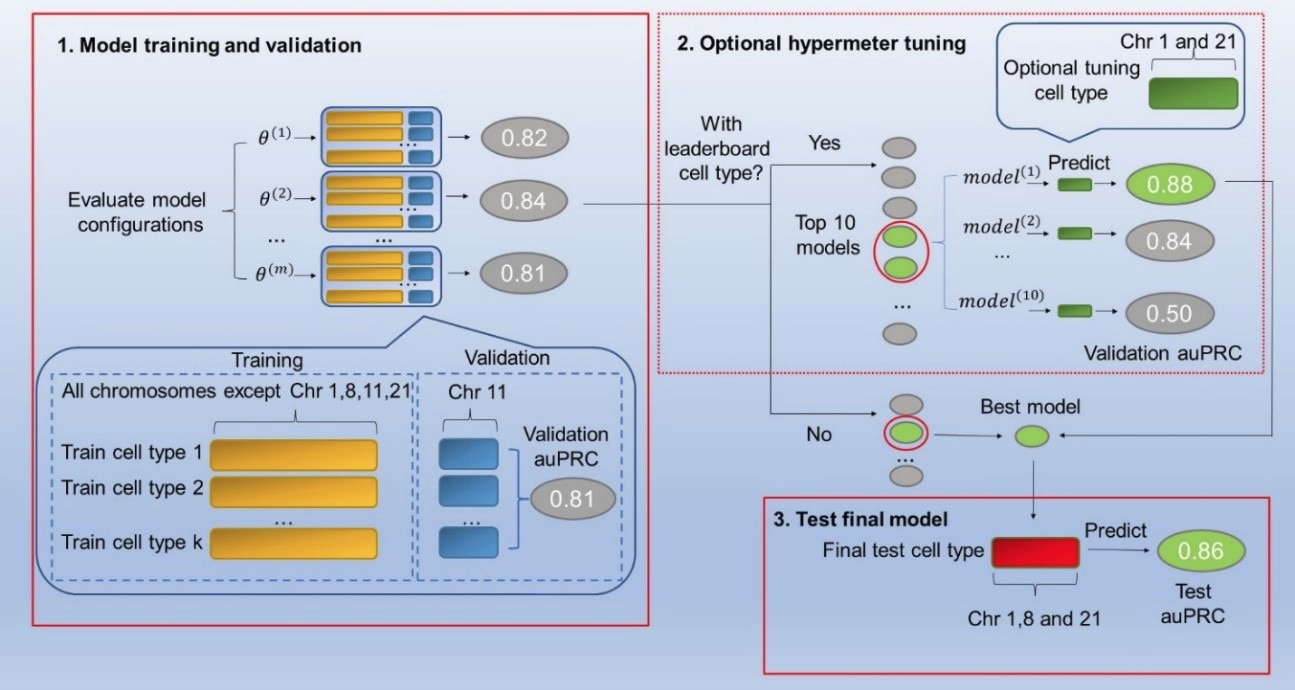
